# A multi-class gene classifier for SARS-CoV-2 variants based on convolutional neural network

**DOI:** 10.1101/2021.11.22.469492

**Authors:** Junyu Fan, Chutao Chen, Chen Song, Jiajie Pan, Guifu Wu

## Abstract

Surveillance of circulating variants of severe acute respiratory syndrome coronavirus 2 (SARS-CoV-2) is of great importance in controlling the coronavirus disease 2019 (COVID-19) pandemic. We propose an alignment-free in silico approach for classifying SARS-CoV-2 variants based on their genomic sequences. A deep learning model was constructed utilizing a stacked 1-D convolutional neural network and multilayer perceptron (MLP). The pre-processed genomic sequencing data of the four SARS-CoV-2 variants were first fed to three stacked convolution-pooling nets to extract local linkage patterns in the sequences. Then a 2-layer MLP was used to compute the correlations between the input and output. Finally, a logistic regression model transformed the output and returned the probability values. Learning curves and stratified 10-fold cross-validation showed that the proposed classifier enables robust variant classification. External validation of the classifier showed an accuracy of 0.9962, precision of 0.9963, recall of 0.9963 and F1 score of 0.9962, outperforming other machine learning methods, including logistic regression, K-nearest neighbor, support vector machine, and random forest. By comparing our model with an MLP model without the convolution-pooling network, we demonstrate the essential role of convolution in extracting viral variant features. Thus, our results indicate that the proposed convolution-based multi-class gene classifier is efficient for the variant classification of SARS-CoV-2.

## 1 Introduction

The Coronavirus disease 2019 (COVID-19) has greatly impacted global public health and all aspects of social activities. According to World Health Organization (WHO) reports, more than 246 million people were diagnosed with COVID-19 as of 31 October 2021, of whom nearly 5 million have died (WHO, 2021). The outbreak is caused by a highly transmissible coronavirus designated as severe acute respiratory syndrome coronavirus 2 (SARS-CoV-2).

Like SARS-CoV and the Middle East respiratory syndrome coronavirus (MERS-CoV), SARS-CoV-2 is a positive-sense RNA virus (Zhou et al., 2020), with a relatively higher mutation rate than double-stranded viruses (Peck and Lauring, 2018). Genetic changes may impact virus phenotypes, such as transmissibility, infectivity, pathogenicity and antigenicity (Harvey et al., 2021), providing the viral population a greater adaptive ability to various environmental conditions. For example, the spike protein substitution D614G enhances SARS-CoV-2 replication, affects susceptibility to antibody neutralization, and has become dominant during the pandemic (Plante et al., 2021). Thus, the emergence and spread of such mutations are problematic concerning both public health and clinical treatment.

Surveillance of circulating SARS-CoV-2 variants benefits vaccine development, development of diagnostics and treatment, and the timely detection of new high-risk variants. Tracking specific variants helps monitoring transmission patterns and adjust public interventions to prevent rapid spread (Meredith et al., 2020; du Plessis et al., 2021). Due to technological improvement in sequencing methods, the reduced per-base cost and sample-to-result turnaround time allow virus genomic sequencing to be widely applied. Unprecedented large-scale viral genomic data have been generated by whole genome sequencing, especially in regions with high sequencing capacity (Oude Munnink et al., 2021). The Basic local alignment search tool (BLAST) (Altschul et al., 1990) is a widely used tool for identifying genomic sequences. Because BLAST compares the query nucleotide sequence against a sequence library, its time consumption increases with the database size. Moreover, BLAST is not effective for bulk data processing. Concerning the rapid increase in the volume of generated genome data, machine learning methods have great potential in the variant classification of SARS-CoV-2. However, raw genomic sequences have no explicit features and are always high-dimensional; therefore, they are poorly depicted by most computational feature selection methods (Angermueller et al., 2016). There is a growing intertest in convolutional neural networks (CNNs), containing multiple non-linear transforming layers, which have been proved useful in extracting significant features from higher-dimensional data (Al-Ajlan and El Allali, 2019) and detecting gene mutation including a single nucleotide polymorphism, an indel variant, and copy number variations (Zhang et al., 2019).

Researchers have done a lot of work on developing novel systems to support COVID-19 diagnosis with the help of CNNs. Marques et al. proposed a CNN model using EfficientNet to detect COVID. The dataset used included 500, 504, and 504 X-ray images from normal, pneumonia, and COVID-19 patients, respectively. Average accuracy for multi-class classification was 96.70% (Marques et al., 2020). El-dosuky et al. presented a cockroach-optimized CNN model to differentiate the DNA sequences of SARS-COV-2 and influenza virus types A, B and C. 65 genome sequences were used for SARS-COV-2 class and 287, 235, and 7 were used for influenza virus type A, B and C, respectively. The classifier achieved an accuracy of 99% (El-Dosuky et al., 2021). However, to the best of our knowledge, no similar study has used CNN methods to classify different variants of SARS-CoV-2. In this study, we propose an alignment-free multi-class classifier based on CNN for four SARS-CoV-2 variants, which may contribute to the methodology of viral variant classification and assist in COVID-19 control.

## 2 Materials and methods

### 2.1 Dataset

The SARS-CoV-2 genomic sequences were collected from the NCBI SARS-CoV-2 Resources (SARS-CoV-2-Sequences, RRID: SCR_018319). This dataset contains sequence profiles of SARS-CoV-2 variants. We collected 2,000 full genomic sequences for each of the four different variants (8,000 genomes in total) with their variant information, including B.1.1.7, B.1.427, B.1.526 and P.1. Each genome is approximately 29,000 base pairs (i.e. letters out of {A, C, G, T, N, T, K, W, Y, R, M, S}). We removed segments where missing values occurred and used the label encoding method to encode the mix-bases symbol gene sequences.

As a common data pre-processing method in machine learning, the collected dataset was randomly split into three separate subsets in an 8:1:1 ratio for training, validation and testing, respectively.

### 2.2 Convolution network

We employed convolution networks to learn the underlying correlation and patterns in the sequence data (Al-Ajlan and El Allali, 2019). Convolutional neurons processes data only for their receptive field, allowing them to capture the local patterns in the genomic sequences.

Considering the discreteness of the genomic sequence data, for the *i*^*th*^ input *I*, its convolution operations is defined as:

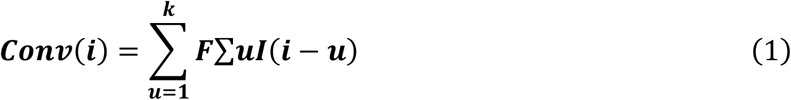

where *F* is the 1-D convolutional filter with a filter size *k* (normally an odd number). Each convolutional layer comprises *n* convolutional filters, transforming the input by arranging the neurons in *n* dimensions. Each convolutional filter has a depth *D*, which is equal to the input depth. The *m*^*th*^ filter ***F***_*m*_ produces a feature map as in Eq (2):

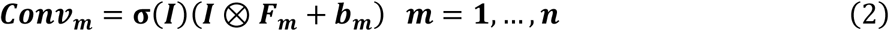

where *σ* is a non-linear activation function (e.g. ReLU (Dahl et al., 2013), sigmoid (Langer, 2021), etc.), ⊗ represents the convolution symbol in Eq (1) and *b*_*m*_ is the bias.

### 2.3 Multilayer perceptron

Multilayer perceptron (MLP) is a fully connected feed-forward artificial neural network that assigns the input to the output through hidden layers, in which the neurons are operated using non-linear activation functions. Typically, three types of activation functions are used: ReLU, sigmoid and tanh.

### 2.4 Logistic regression

Since the output for variant classification is categorical, we use logistic regression as the output layer of our classifier to generate the results. For input feature *x*_*i*_ and label *y*_*i*_, the logistic regression model can be presented as shown in Eq (3) and Eq (4):

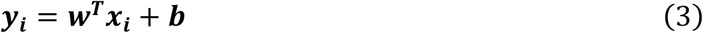

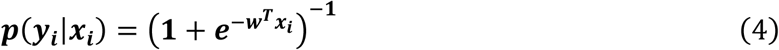

where the weight matrix *w* and the bias *b* are trained to minimize the objective function.

### 2.5 Loss function (cross-entropy)

The loss function represents the difference between the label *y* and predicted result ŷ. Different loss functions could be used in a machine learning model. The cross-entropy objective was used for our classifier:

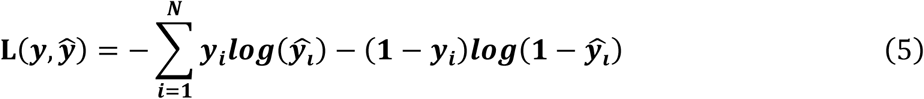

### 2.6 Convolution-based gene classifier

To build a predictive model for classifying virus variants, we constructed a deep learning model utilizing a stacked 1-D convolutional neural network and MLP. As shown in Figure 1. and pipeline (Figure 2), the proposed model comprises ten layers, three stacked convolution-pooling nets, 2-layer MLP, one input, and one output. The input layer takes the input as the labeled gene sequence data. The output layer of the classifier is a logistic regression model that generates the probabilities of each type of variant. Stacked convolution-pooling nets are used as hidden layers, followed by MLP layers. The model was trained and optimized using a backpropagation algorithm with an Adam optimizer. We determined the hyperparameters using the grid research method. We used cross-validation to select the optimal model and evaluated model performance on an independent dataset.

**Figure 1.**
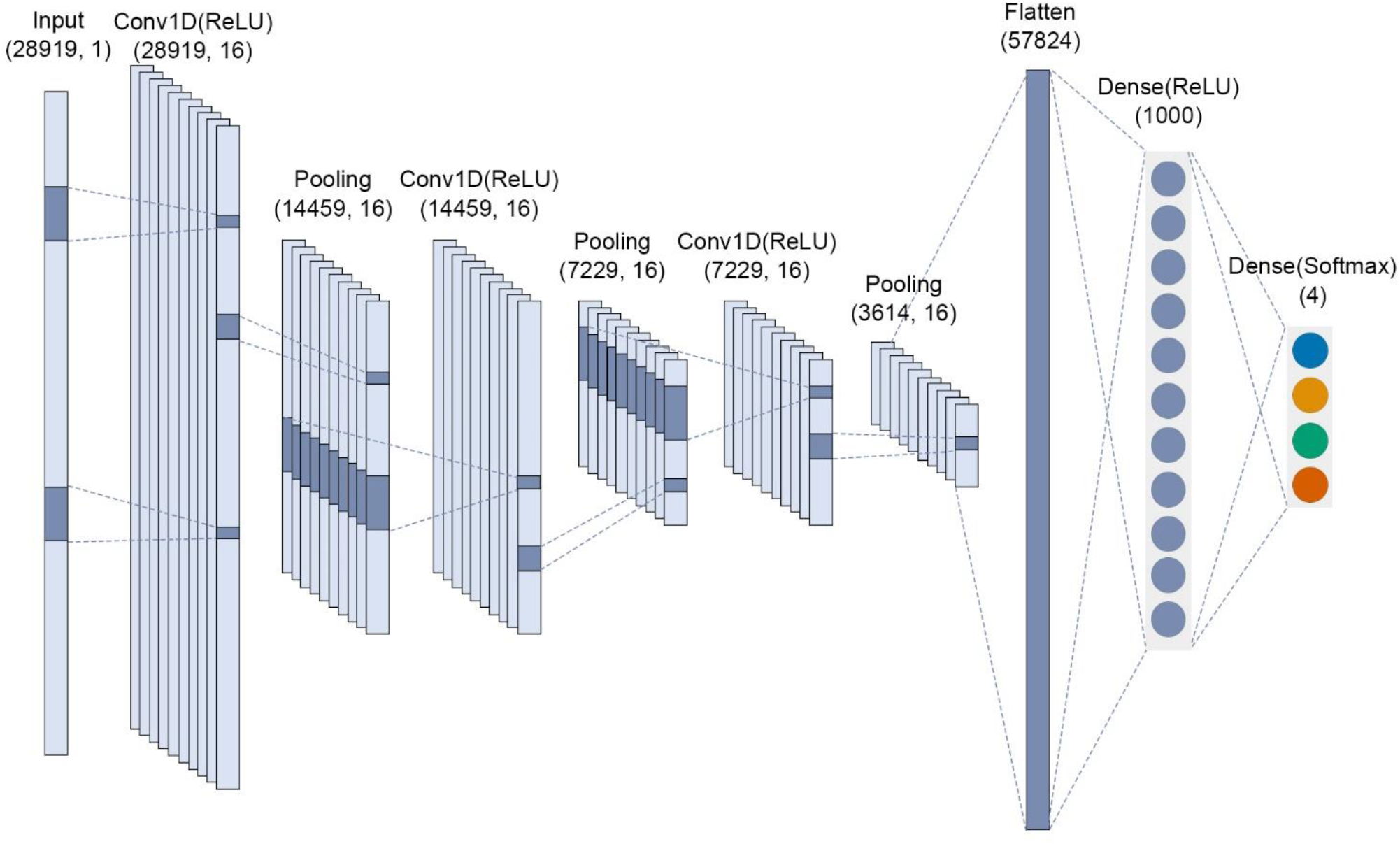
The architecture of the proposed CNN multi-class gene classifier.

**Figure 2.**
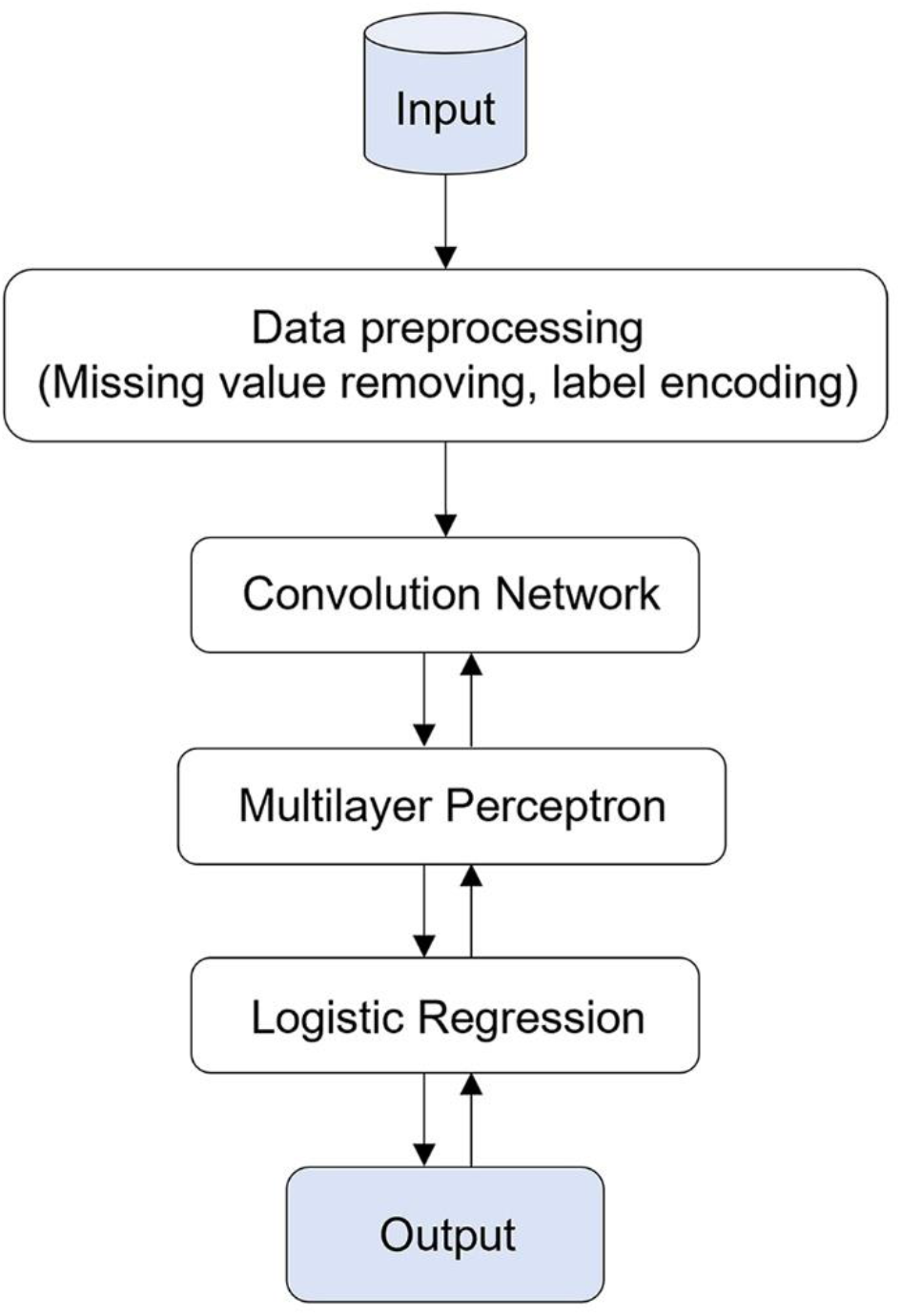
Pipeline of the proposed work. ↑: parameter training/backpropagation/optimization

### 2.7 Implementation and evaluation

The experiments were carried out on Python 3.8 on an Intel® CoreTM i7, 2.00 GHz with 8GM RAM. Logistic regression (LR), K-nearest neighbor (KNN), support vector machine (SVM) and random forest (RF) are implemented with Scikit-learn (Pedregosa et al., 2011) while MLP and CNN were implemented with TensorFlow (Abadi et al., 2016). The codes related are available at https://github.com/chotiu5/CNN.

The metrics used to evaluate the proposed model include accuracy, precision, recall and F1 score, which are calculated according to the number of true positives (TP), true negatives (TN), false positives (FP) and false negatives (FN):

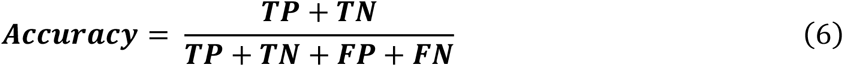

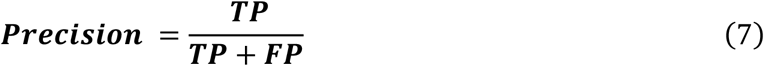

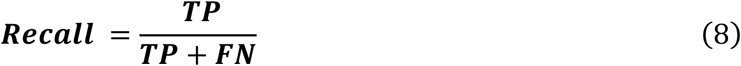

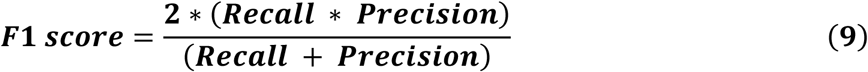

The macro-averages of precision, recall and F1 scores were used in this study in consideration of the balanced dataset. The macro-average scores were calculated as the arithmetic mean of individual classes’ corresponding scores.

In the external validation, a confusion matrix, also called an error matrix, was used to visualize the predictive performance of the model in the testing data by showing the number of correct and incorrect predictions for each viral variant. t-distributed stochastic neighbor embedding (t-SNE) is a widely used statistical method for converting the distance of high-dimensional data into conditional probabilities representing similarities (Van der Maaten and Hinton, 2008). We used t-SNE to visualize the separability of different variants before and after training with the value of perplexity set as 38. The predictive ability of this multi-class classifier for testing data is also illustrated by receiver operating characteristic (ROC) curves and the area under the ROC curve (AUC).

## 3 Results and discussion

This paper proposes an in silico approach for classifying SARS-COV-2 variants based on their genomic sequences. We first used the learning curves of the CNN gene classifier to observe the training process of our CNN gene classifier. Next, the proposed classifier was evaluated using stratified 10-fold cross-validation. We then used external validation and visualization to test the performance of the classifier. Finally, we compared the performance of the proposed CNN classifier with that of other machine learning methods to further validate the ability of the classifier.

### 3.1 Training process of the CNN gene classifier

Loss curves and accuracy curves are shown for 70 epochs to observe the performance of our CNN gene classifier during training. As shown in Figure 3, the initial loss value and accuracy of validation were 0.2511 and 0.9050, respectively. The classifier had fast training speed. The validation accuracy exceeded 0.9675 and the validation loss dropped down to 0.1107 after two epochs. As the epoch count reached 23, the classifier achieved a validation loss of 0.0095 and validation accuracy of 0.9987. When the epoch number increases to more than 23, the accuracy and loss values tend to be stable. The training and validation values are almost identical, indicating that the classifier has outstanding generalization capability without distinct overfitting.

**Figure 3.**
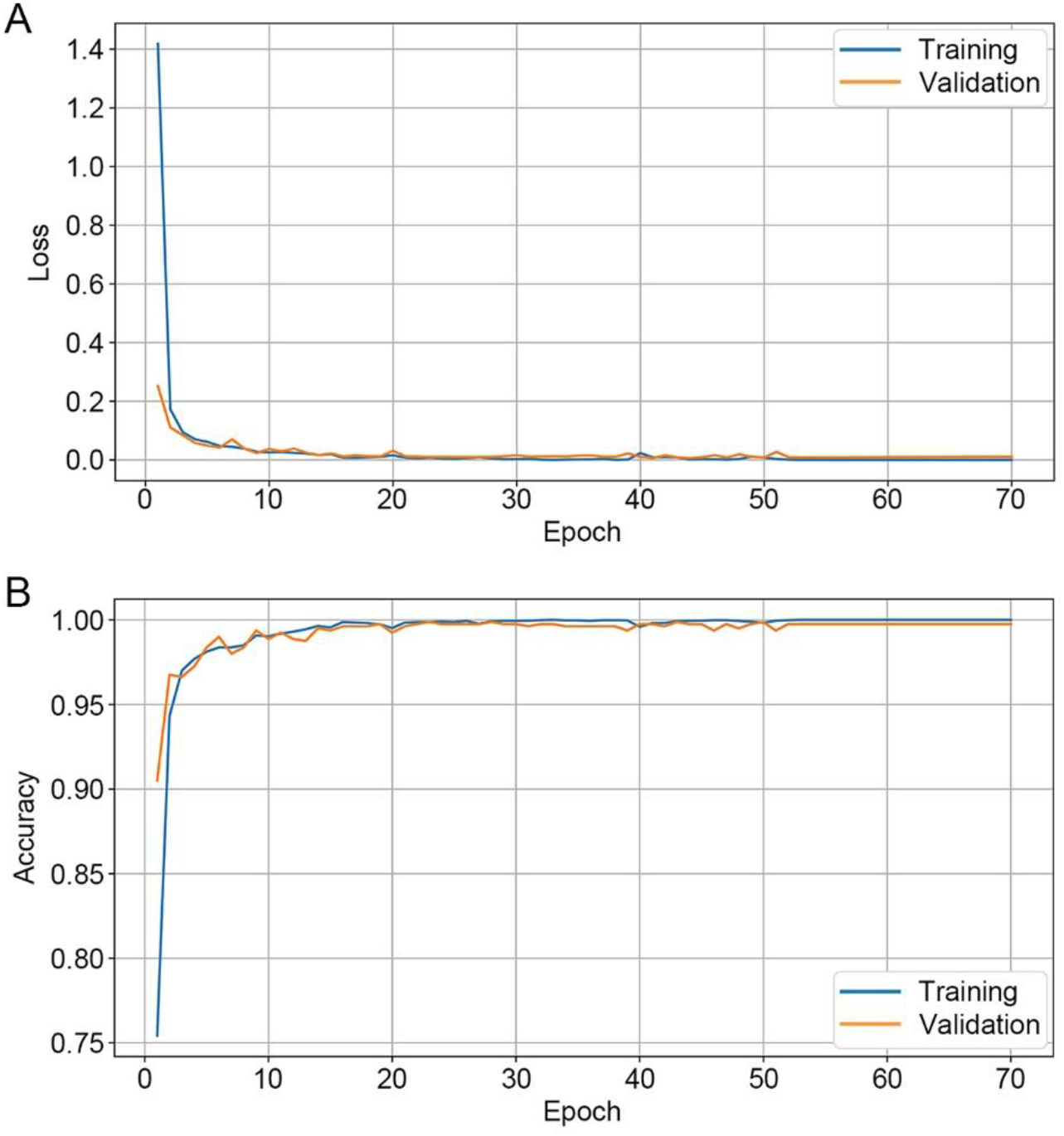
Learning curves of the multi-class CNN gene classifier. **(A)** Loss curves for training data and validation data. **(B)** Accuracy curves for training data and validation data.

### 3.2 Stratified 10-fold validation of the CNN gene classifier

The stratified 10-fold cross-validation method was applied to a joined dataset of the training and validation data for unbiased evaluation of multiple train-test splits. The joined dataset was shuffled and split into ten groups containing the same proportions of the four variants. Nine groups were used to train the model within each fold, while the remaining 1 group was assigned for validation. The performance metrics obtained in each fold are listed in Table 1. The highest accuracy, precision, recall and F1 score 0.9986 were reported at the 5th, 6th and 10th fold, while the lowest values of 0.9917 were observed at the 3rd and the 4th folds. The average results of all folds were 0.9949, indicating the good prediction ability of the proposed model.

**Table 1.** Classification results using stratified 10-fold cross-validation.

| Fold | Accuracy | Precision | Recall | F1 Score |
| --- | --- | --- | --- | --- |
| 1 | 0.9944 | 0.9945 | 0.9944 | 0.9944 |
| 2 | 0.9931 | 0.9931 | 0.9931 | 0.9931 |
| 3 | 0.9917 | 0.9917 | 0.9917 | 0.9917 |
| 4 | 0.9917 | 0.9918 | 0.9917 | 0.9917 |
| 5 | **0.9986** | **0.9986** | **0.9986** | **0.9986** |
| 6 | **0.9986** | **0.9986** | **0.9986** | **0.9986** |
| 7 | 0.9958 | 0.9958 | 0.9958 | 0.9958 |
| 8 | 0.9931 | 0.9931 | 0.9931 | 0.9931 |
| 9 | 0.9931 | 0.9931 | 0.9931 | 0.9931 |
| 10 | **0.9986** | **0.9986** | **0.9986** | **0.9986** |
| Average | 0.9949 | 0.9949 | 0.9949 | 0.9949 |

The best results for each evaluation indicator are highlighted in bold.

### 3.3 External validation and visualization of CNN classifier performance

The 10% holdout subset of the original dataset is then applied for the final estimation to ensure that the CNN gene classifier can generalize well to new, unseen data. The accuracy, precision, recall and F1 score for each variant are shown in Table 2. There is only little difference in the predictive performance of the proposed classifier applying to different variants. The overall accuracy for multi-class classification is 0.9962. Figure 4 shows the ROC curves of the classifier. They further proved that the CNN classifier performed well for all the variants as the area under the ROC curve (AUC) is approximately 1.00. Figure 5 shows the confusion matrix of the classifier used to visualize the correct and incorrect classifications. This indicates that the proposed classifier has a low rate of error identification.

**Table 2.** Variant classification results of external validation.

| Variant | Accuracy | Precision | Recall | F-score |
| --- | --- | --- | --- | --- |
| B.1.1.7 | NA | 1.0000 | 0.9950 | 0.9975 |
| B.1.427 | NA | 0.9950 | 1.0000 | 0.9975 |
| B.1.526 | NA | 0.9901 | 1.0000 | 0.9950 |
| P.1 | NA | 1.0000 | 0.9900 | 0.9950 |
| All | 0.9962 | 0.9963 | 0.9963 | 0.9962 |

NA, not applicable.

**Figure 4.**
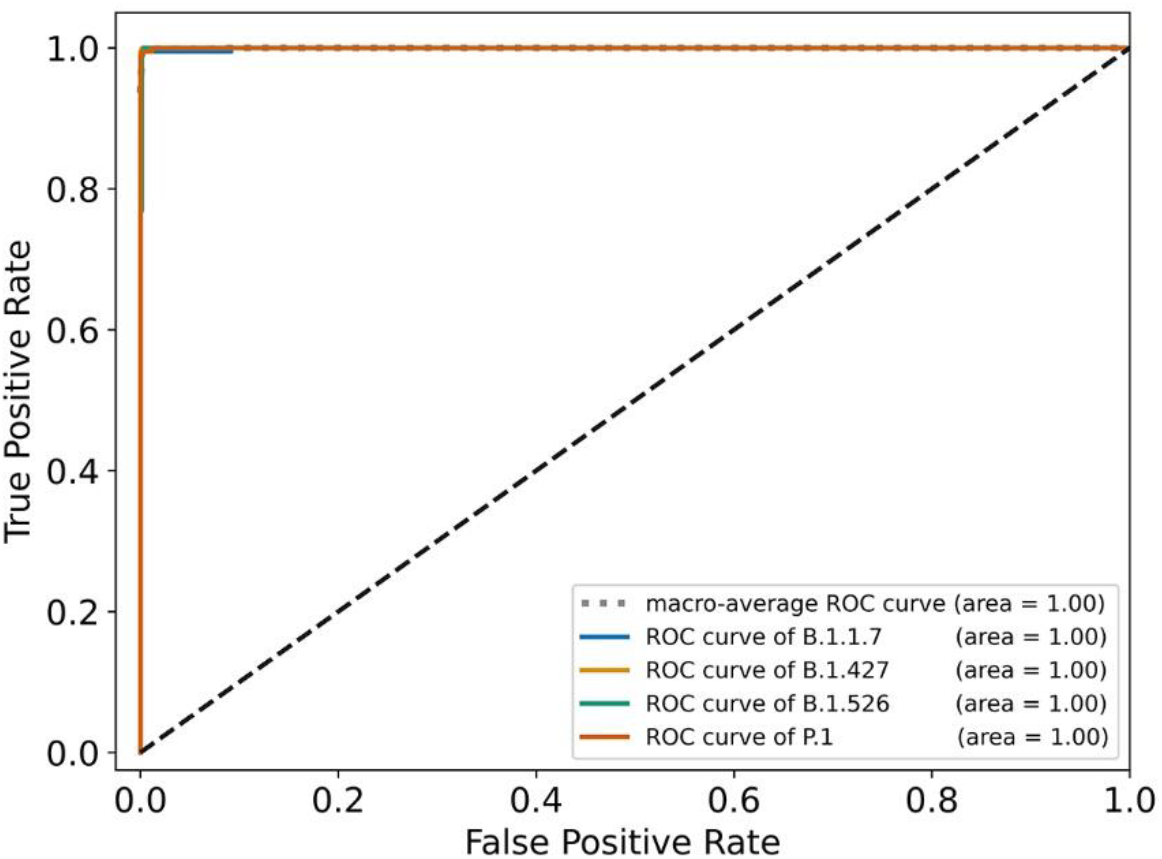
The ROC curves of external validation for four specific viral variants and their macro-average ROC curve.

**Figure 5.**
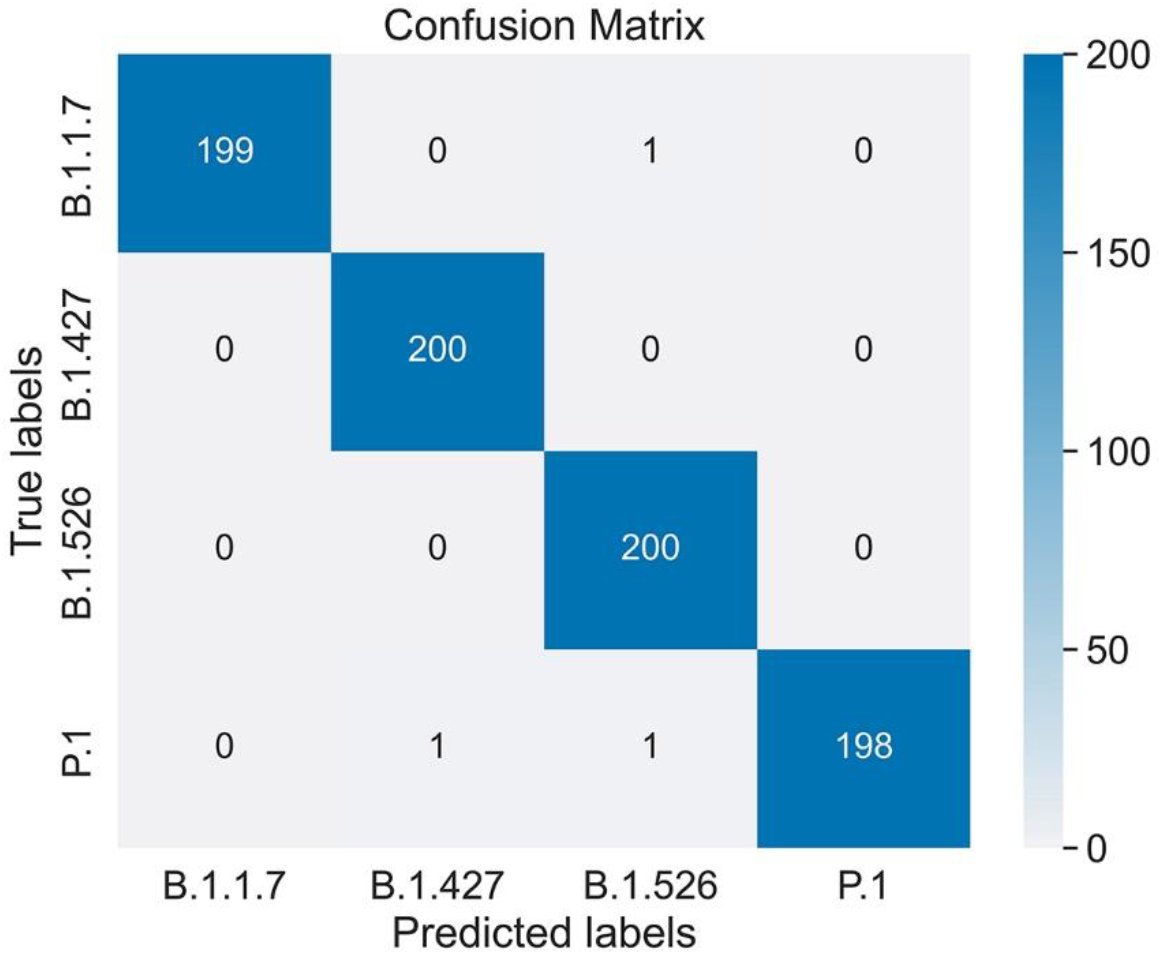
Confusion matrix of external validation for the CNN multi-class gene classifier.

The performance of our CNN classifier was also visualized using t-SNE. The two-dimensional maps generated from the raw SARS-CoV-2 genome data showed strong overlap between variants and did not appear to be easily separable (Figure 6A, 6C). After being processed by the trained CNN classifier, the separation visibility among variant clusters was greatly improved both in the training and testing data (Figure 6B, 6D). This result indicates that our proposed model can effectively extract features from the four viral variants for classification.

**Figure 6.**
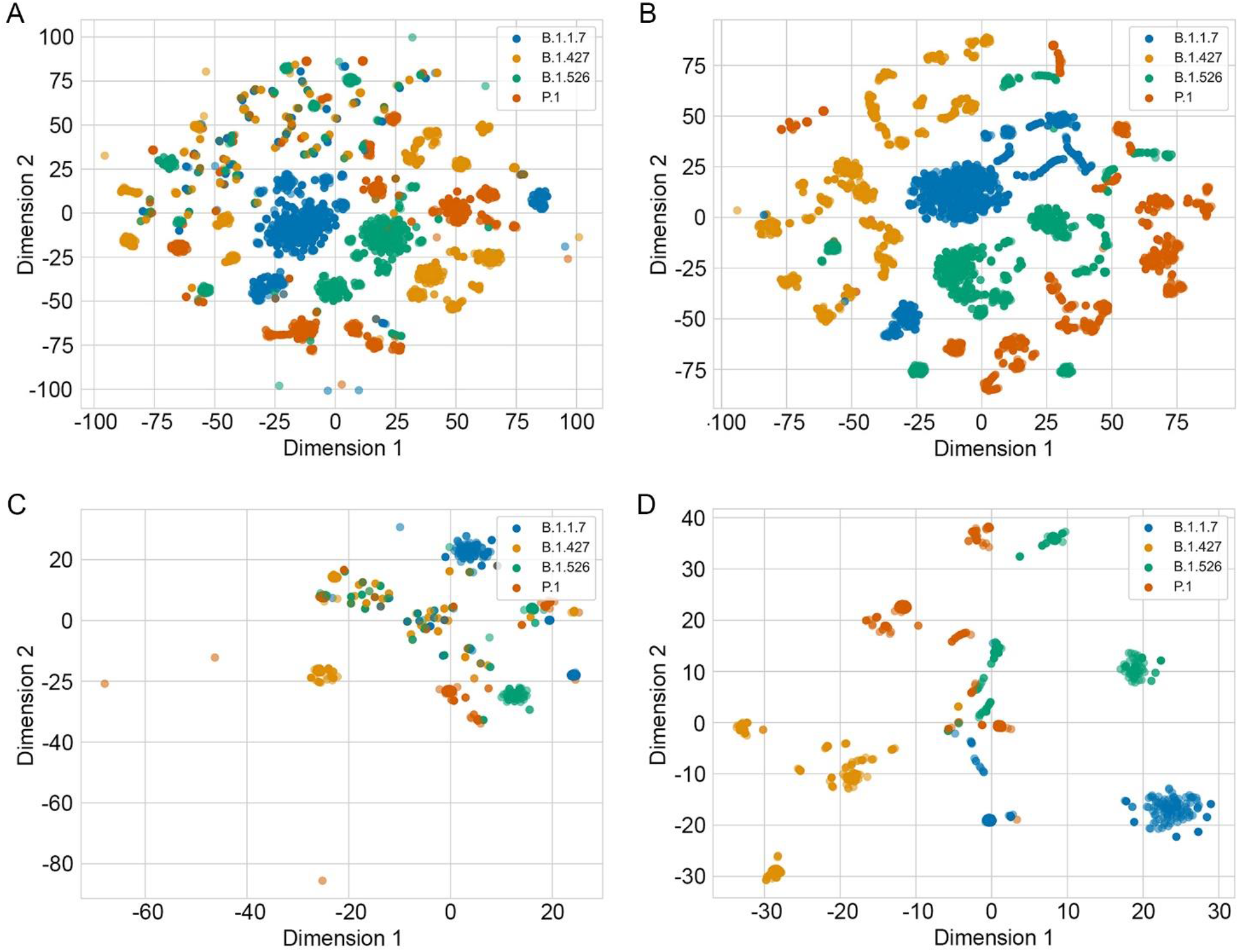
t-SNE two-dimensional visualization of SARS-CoV-2 genome data set. **(A)** t-SNE distribution of the raw training and validation data. **(B)** t-SNE distribution of the training and validation data after being processed by the classifier. **(C)** t-SNE distribution of the raw external testing data. **(D)** t-SNE distribution of the testing data after being processed by the classifier.

### 3.4 Compared to other methods

Finally, as shown in Table 3, the CNN classifier performs better than other machine learning methods including LR, KNN, SVM, and RF. Notably, a neural network without the convolutional layers, namely MLP, performs worse than the other methods. The training loss continued to decrease after 25 epochs of training, whereas the validation loss increased, indicating over-fitting of the MLP model. The accuracy values show large fluctuations and cannot converge at a high level in 200 epochs (Figure 7). In contrast, the CNN classifier preformed the highest best in both stratified 10-fold validation and holdout dataset testing. Thus, it is demonstrated that the convolutional layers are of great importance in extracting features from the genomic sequences of the four viral variants. This can be interpreted based on the biological characteristics of the genome. Although the genome is primarily structured with nucleotides, it contains more information than just isolated bases and their corresponding loci. The nucleotides conduct downstream phenotype of an organism in the form of continuous fragments interacting with their “neighbors”, such as codons, coding regions, cis-acting elements, transcription-regulating sequences (TRSs), splice sites, etc. (Sola et al., 2015; Cross et al., 2019). Convolution layers allow the local input to be spatially aggregated so that the model can learn through the complicated composition of genomic sequences (LeCun et al., 2015), some of which may be unexplainable under the current stage of biological knowledge (Zrimec et al., 2020).

**Table 3.** Comparative performance of classifiers based on CNN or other machine learning methods.

| Model | 10-fold validation |  |  |  | External validation |  |  |  |
| --- | --- | --- | --- | --- | --- | --- | --- | --- |
|  | Accuracy | Precision | Recall | F1 Score | Accuracy | Precision | Recall | F1 Score |
| LR | 0.9904 | 0.9905 | 0.9904 | 0.9904 | 0.9888 | 0.9888 | 0.9887 | 0.9887 |
| KNN | 0.9771 | 0.9776 | 0.9771 | 0.9771 | 0.9838 | 0.9838 | 0.9838 | 0.9837 |
| SVM | 0.9633 | 0.9643 | 0.9633 | 0.9632 | 0.9600 | 0.9607 | 0.9600 | 0.9598 |
| RF | 0.9922 | 0.9922 | 0.9922 | 0.9922 | 0.9925 | 0.9925 | 0.9925 | 0.9925 |
| MLP | 0.9197 | 0.9316 | 0.9197 | 0.9187 | 0.9550 | 0.9578 | 0.9550 | 0.9550 |
| CNN | 0.9949 | 0.9949 | 0.9949 | 0.9949 | 0.9962 | 0.9963 | 0.9963 | 0.9962 |

LR, logistic regression; KNN, K-nearest neighbor; SVM, support vector machine; RF, random forest; MLP, multilayer perceptron; CNN, convolutional neural network.

**Figure 7.**
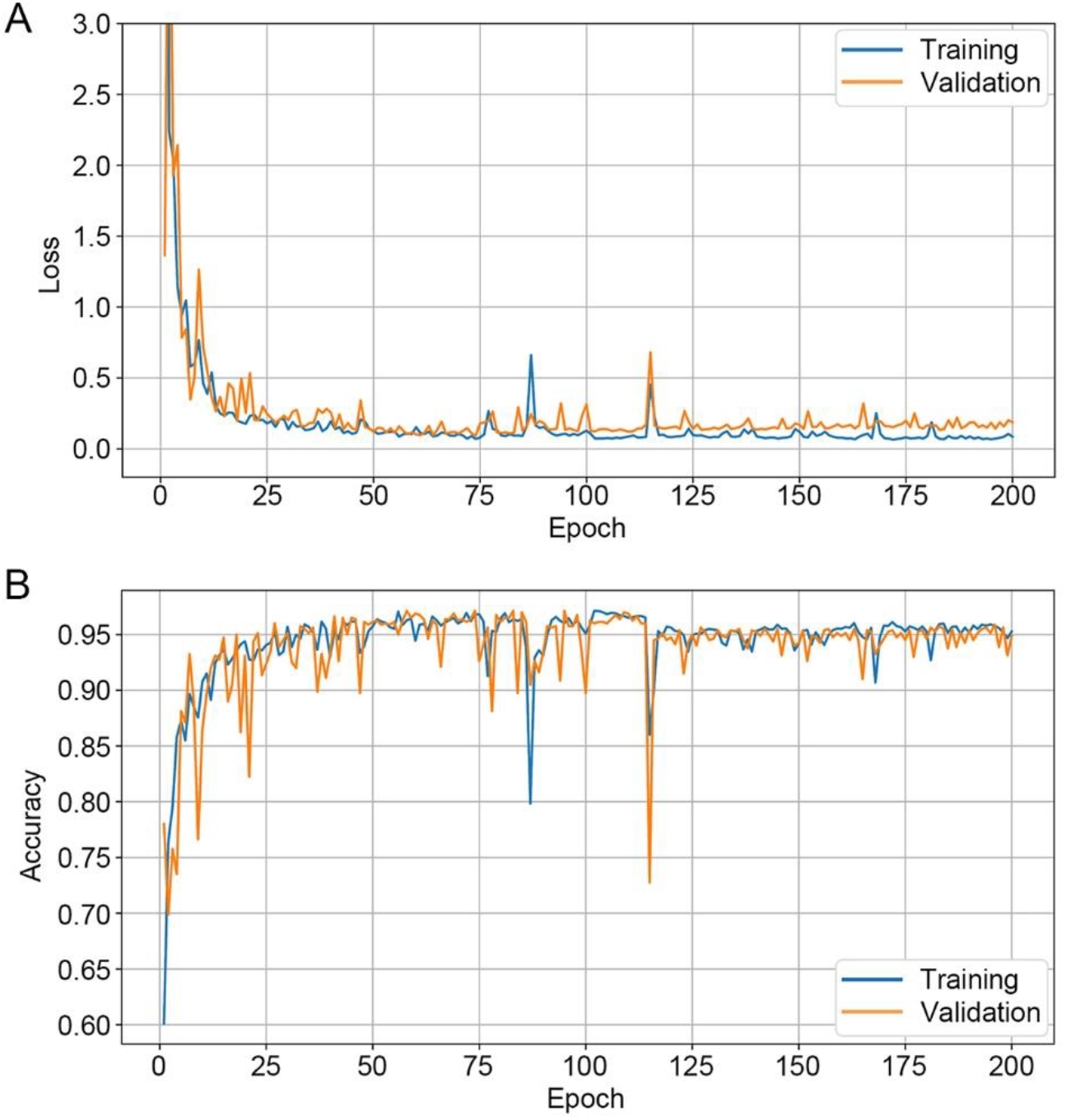
Learning curves of MLP. **(A)** Loss curves for training and validation data. **(B)** Accuracy curves for training and validation data.

## 4 Conclusion

This study reports an automated method based on CNN for SARS-CoV-2 variant classification with an accuracy of 0.9962. By visualizing the learning process and data separability, and comparing it with other machine learning methods, we demonstrated that the proposed CNN gene classifier could effectively extract features from the collected four variants of SARS-CoV-2 genomic data and enable robust variant classification.

## Acknowledgments

We would like to express our gratitude to all the frontline healthcare workers for their dedication to human health during the COVID-19 pandemic.

## Data Availability Statement

The datasets analyzed in this study are available at https://github.com/chotiu5/CNN.

